# Tenotomy-induced muscle atrophy is sex-specific and independent of NFκB

**DOI:** 10.1101/2022.07.26.501541

**Authors:** Gretchen A. Meyer, Stavros Thomopoulos, Yousef Abu-Amer, Karen C. Shen

**Affiliations:** Program in Physical Therapy, Washington University School of Medicine, St. Louis, MO, USA; Departments of Neurology and Biomedical Engineering, Washington University School of Medicine, St. Louis, MO, USA; Department of Orthopaedic Surgery, Washington University School of Medicine, St. Louis, MO, USA; Departments of Orthopaedic Surgery and Biomedical Engineering, Columbia University, New York, NY, USA; Department of Cell Biology & Physiology, Washington University School of Medicine, St. Louis, MO, USA; Shriners Hospital for Children, St. Louis, MO, USA

**Keywords:** Rotator Cuff, IKKβ, Muscle Wasting

## Abstract

The nuclear factor-κB (NFκB) pathway is a major thoroughfare for skeletal muscle atrophy and is driven by diverse stimuli. Targeted inhibition of NFκB through its canonical mediator IKKβ effectively mitigates loss of muscle mass across many conditions, from denervation to unloading to cancer. In this study, we used gain- and loss-of-function mouse models to examine the role of NFκB in muscle atrophy following rotator cuff tenotomy – a model of chronic rotator cuff tear. IKKβ was knocked down or constitutively activated in muscle-specific inducible transgenic mice to elicit a 2-fold gain or loss of NFκB signaling. Surprisingly, neither knockdown of IKKβ nor overexpression of caIKKβ significantly altered the loss of muscle mass following tenotomy. This finding was consistent across measures of architectural adaptation (fiber cross-sectional area, fiber length, fiber number), tissue pathology (fibrosis and fatty infiltration) and intracellular signaling (ubiquitin-proteasome, autophagy). Intriguingly, late-stage tenotomy-induced atrophy was exacerbated in male mice compared to female mice. This sex specificity was driven by ongoing decreases in fiber cross-sectional area, which paralleled the accumulation of large autophagic vesicles in male, but not female muscle. These findings suggest that tenotomy-induced atrophy is not dependent on NFκB and instead may be regulated by autophagy in a sex-specific manner.

## 1 Introduction

At the most basic level, skeletal muscle atrophy is a shift in the physiological balance between protein synthesis and breakdown. This shift can be initiated by a wide variety of stimuli, from disuse to starvation to cancer, but all eventually converge on a common result – reduced muscle mass. As expected from the diverse stimuli, atrophy can be initiated through a variety of intracellular pathways. However, studies have found surprising commonality downstream, leading to identification of a number of “lynchpin” mediators – in particular signaling through nuclear factor kappa beta (NFκB), which has been described as both necessary and sufficient for skeletal muscle atrophy (Jackman *et al*., 2013; Sandri, 2013).

Our current understanding of NFκB’s role in adult muscle atrophy has been reviewed in detail (Jackman *et al*., 2013). In brief, NFκB is a family of transcription factor subunits that exist as dimers and are localized to the cytoplasm by inhibitory kinases (inhibitor of κB, IκB). Upon stimulation, IκB kinases (IKKα and IKKβ) are phosphorylated and in turn phosphorylate IκB (predominantly IκBα). This leads to cytosolic degradation of Iκβ and nuclear transport of NFκB transcription factor dimers to initiate transcription of genes involved in the atrophy program. Inhibiting IKKβ activity by muscle specific knockout of its gene (*Ikbkb*) or electroporating a dominant negative form into skeletal muscle prevented approximately half the loss of muscle mass induced by denervation (Mourkioti *et al*., 2006), unloading (Van Gammeren *et al*., 2009), or immobilization (Reed *et al*., 2011). Similar results were found in the Iκβα superrepressor mouse, which displays dampened NFkB activity, in response to an inflammatory insult (Haegens *et al*., 2012; Langen *et al*., 2012) denervation (Cai *et al*., 2004), unloading (Judge *et al*., 2007), nutrient deprivation (Lee & Goldberg, 2015) or tumor (Cai *et al*., 2004). Assessment of NFκB activity by nuclear localization or DNA binding of its subunits confirmed that inhibition of IKKβ activity or IκBα degradation completely (Cai *et al*., 2004; Senf *et al*., 2008; Van Gammeren *et al*., 2009) or partially (Mourkioti *et al*., 2006) blocked NFκB translocation, demonstrating that NFκB signaling is responsible for a considerable portion of atrophy driven by these diverse stimuli.

The best studied gene targets of NFκB in muscle are the E3 ubiquitin ligases muscle ring finger 1 (MuRF1) and muscle atrophy F-box (MAFbx; a.k.a. Atrogin-1) (Wu *et al*., 2011). These genes, termed “atrogenes”, are thought to control the majority of proteolysis in skeletal muscle across atrophic conditions (Bodine *et al*., 2001; Gomes *et al*., 2001). Knockout of MuRF1 results in a similar extent of muscle sparing to NFκB inhibition in denervation (Gomes *et al*., 2012), unloading (Labeit *et al*., 2010), glucocorticoid treatment (Baehr *et al*., 2011) and inflammatory insult (Files *et al*., 2012) leading it, too, to be labeled “essential” for skeletal muscle atrophy (Peris-Moreno *et al*., 2020). While compelling, this explanation remains incomplete. First, because inhibition of NFκB or MuRF1 signaling does not completely prevent atrophy induced by these various stimuli and second because the degree of prevention varies by stimulus and muscle. For example, muscle specific *Ikbkb* knockout prevented ∼70% of denervation-induced decrease in mass of the soleus muscle, but only ∼30% in the gastrocnemius muscle (Mourkioti *et al*., 2006). Similarly, Iκβα super-repressor prevented only ∼25% of the denervation-induced mass loss in the gastrocnemius and tibialis anterior muscles but ∼50% of the tumor-induced loss in mass (Cai *et al*., 2004).

Recently, tendon tears (leading to release of muscle tension and muscle retraction) have emerged as potentially employing distinctive mechanisms of muscle atrophy. A comparative study of atrophic signaling between tenotomy and immobilization found that tenotomized gastrocnemius muscle atrophy was primarily characterized by a lysosomal/autophagic signature, in contrast to the classical ubiquitin-mediated proteasomal activity signature of the immobilized gastroc (Bialek *et al*., 2011). MuRF1 or MAFbx expression did not significantly increase with tenotomy, where dramatic increases were again shown with immobilization, denervation, and glucocorticoid-induced atrophy. This is somewhat surprising given the similarity in signaling induced by other models of unloading that remove either passive loading (e.g., hindlimb suspension and immobilization by casting) or active loading (e.g., denervation). In fact, previous to this study, tenotomy was assumed to employ the ubiquitin-mediated proteasome activity through NFκB and the atrogenes simply because of this similarity (Laron *et al*., 2012). More recent studies have focused on tenotomy of the rotator cuff (RC) muscles as a model to investigate the mechanisms of muscle atrophy in human chronic RC tears. These studies have confirmed no increases in MuRF1 or MAFbx following RC tenotomy (Liu *et al*., 2012) and have uncovered additional evidence for increased autophagic flux (Gumucio *et al*., 2012; Joshi *et al*., 2014). Evidence in support of proteasomal activity in the early phase of RC tenotomy, however, also exists (Valencia *et al*., 2017), leaving open the possibility of coordinated action of multiple pathways.

Taken together, these prior studies suggest that although NFκB signaling is a critical component of many disparate atrophy models, it may not be ubiquitous - and in particular may not play a major role in tenotomy-induced atrophy. However, due to the complexity of NFκB signaling, its role in tenotomy-induced atrophy (or lack thereof) cannot be inferred from the lack of atrogene expression or the autophagic state of the muscle alone. First, because evidence suggests that NFκB is capable of driving atrophy independent of MuRF1 or MAFbx (Cai *et al*., 2004) and second, because there is a complex link between NFκB and autophagy which could cause NFκB to drive the autophagic process (reviewed in (Xia *et al*., 2021)). In the current study, we sought to determine the role of NFκB signaling in tenotomy-induced atrophy of the RC muscles using muscle specific gain- and loss-of-function IKKβ transgenic mouse models to inhibit and promote NFκB activity, respectively. We hypothesized that NFκB inhibition would ameliorate tenotomy-induced atrophy through an atrogene-independent mechanism, thus supporting NFκB inhibitors as a novel therapeutic strategy to maintain muscle mass following human chronic rotator cuff tears. However, we found that knockdown of IKKβ had no impact on the loss of muscle mass or contractile performance following tenotomy and that overexpression of constitutively active IKKβ was insufficient to drive it. Instead, our data point to an NFκB-independent, sex-specific mechanism driven by autophagy.

## 2 Results

### Tenotomy-induced atrophy of the rotator cuff muscles is sex-specific

Change in muscle mass was the first outcome assessed following tenotomy of the supraspinatus (SS) and infraspinatus (IS) muscles – the primary abductors in the rotator cuff. Although this study was primarily concerned with differences between genotypes (due to changes in NFκB signaling), differences were noted between males and females in the wildtype group. Specifically, at week 8 (W8) post-tenotomy, mass loss in the male supraspinatus (SS) and infraspinatus (IS) muscles outpaced that in female mice (Fig 1A). This difference resulted in a significant time x sex ANOVA interaction in both muscles over the time course investigated and a significant difference between male and female W8 groups by post-hoc comparison. Further investigation in the other outcomes of the study suggest these sex-specific muscle mass effects arose from sex-specific differences in the architectural mechanisms of atrophy. Atrophy is driven by three primary architectural adaptations: decreases in fiber sizes in the radial dimension (fiber atrophy), decreases in fiber numbers (hypoplasia), and decreases in fiber sizes in the longitudinal dimension (sarcomere subtraction). Only male mice demonstrated decreased fiber cross-sectional area (CSA) at W8 post tenotomy. Analysis of type 2b fiber CSA, the primary fiber type in both SS and IS muscles, showed a significant time x sex ANOVA interaction and a significant difference between male and female W8 groups by post-hoc comparison (Fig 1B). Conversely, only female mice decreased fiber numbers at W8 post tenotomy, with a significant difference between male and female groups by post-hoc comparison (Fig 1C). Female mice also decreased fiber lengths to a greater extent than male mice at both W2 and W8 (Fig 1D).

**Figure 1.**
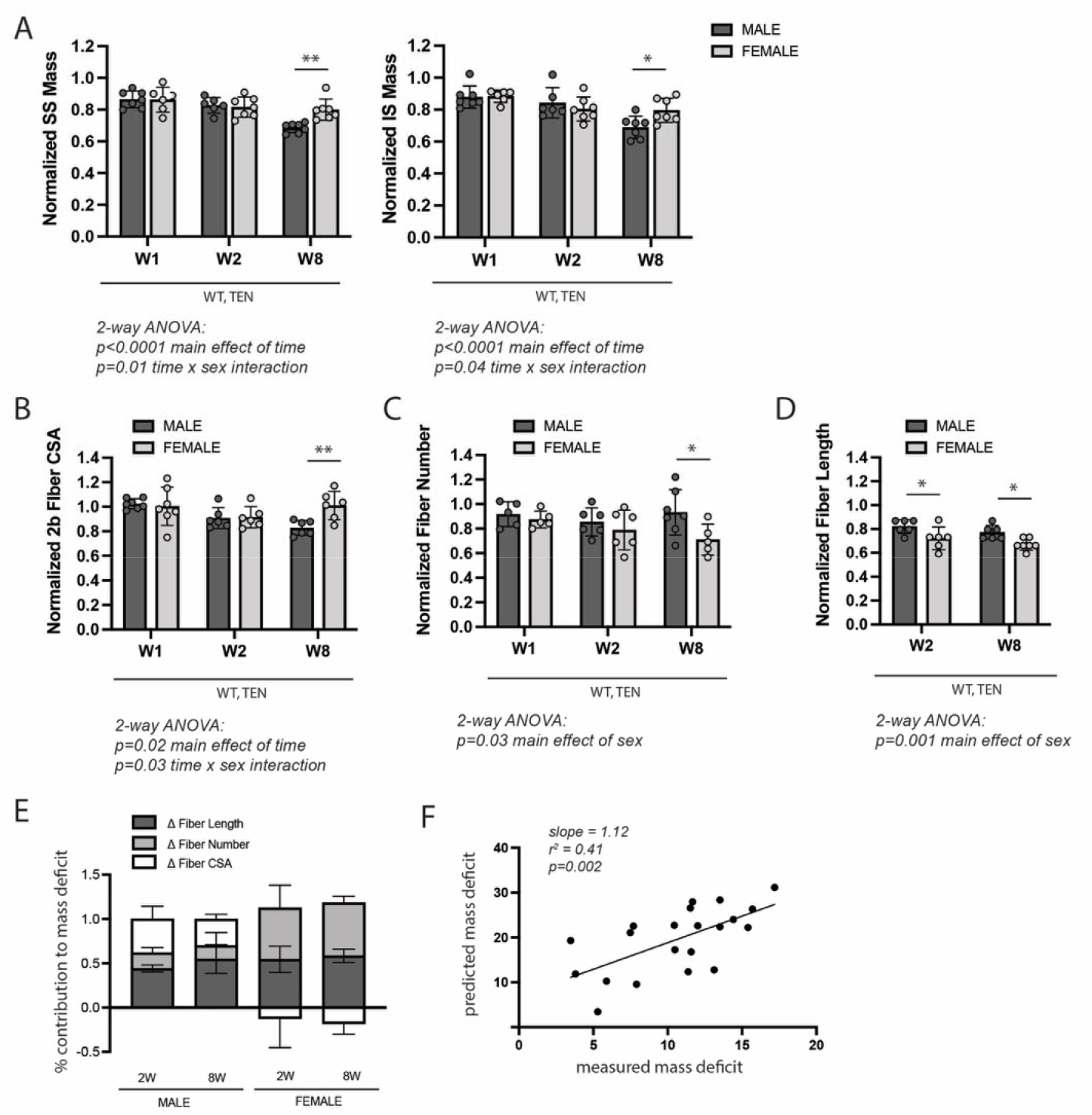
WT control mice exhibited sex-specific responses to late-stage tenotomy-induced muscle atrophy. (A) Supraspinatus (SS) and infraspinats (IS) muscles from male mice lose more mass than female muscles at 8 weeks (W8) post tenotomy (TEN). Muscle mass is normalized to body mass and the average value from the sham group for each sex. (B) Type 2b fiber cross-sectional area (CSA), assessed in histological sections, is also significantly reduced in males compared with females at W8. (C) The total number of fibers counted per histological section is significantly reduced in the female SS compared with male at W8. (D) Prediction of fiber lengths from measured muscle lengths during physiological testing indicates that female SS muscles lose a greater fraction of their fiber length at both W2 and W8 post-tenotomy. (G) Predictions of the contributions of each architectural change (B-D) to the measured mass deficit (A). Change in (Δ) fiber length (dark gray), fiber number (light gray), fiber CSA (white) show different trends for males and females at W2 and W8 post-tenotomy. (F) The mass deficit *predicted* by architectural changes and the mass deficit *measured* at W2 and W8 are significantly correlated. (A-D) Raw values were normalized to the average of the sham group of the same sex. N=5-7 per group; * p<0.05, **p<0.01

To better understand the relative impact of these architectural changes on muscle mass, a geometrical model was employed to predict the mass deficit from each change. Overall, changes in fiber length accounted for most of the mass deficit with tenotomy in both sexes (Fig 1E, purple). However, there were notable differences in the other sources. While changes in fiber CSA accounted for 25-30% of the mass loss in male mice at 2W and 8W, they accounted for none of the mass loss in female mice. In contrast, changes in fiber number account for 30-40% of the mass loss in females, but only 15-20% in males. To validate that our model captured the major sources of mass loss, we regressed the predicted mass deficit against the measured mass deficit for each muscle across both sexes. The measurements were significantly correlated (r2=0.41, p=0.002; Fig 1F). Additionally, the slope of the regression line was close to 1, indicating that the architectural measurements were able to account for the majority of muscle mass loss following tenotomy. Given these differences, all subsequent analyses investigating the effect of IKKβ knockdown and overexpression on tenotomy-induced muscle atrophy consider sex-specific effects.

### IKKβ gain- and loss-of-function did not affect tenotomy-induced muscle atrophy

The gene encoding IKKβ (*Ikbkb*) was selectively deleted (IKKb^MKD^) or constitutively activated (IKKb^MCA^) in mature skeletal muscle fibers of 6-8 month old transgenic mice. Tamoxifen treatment of IKKb^MKD^ mice caused a 50-60% reduction in the expression of *Ikbkb* mRNA in supraspinatus muscles that was sustained for through W8 (Fig 2A). Conversely, tamoxifen treatment of IKKb^MCA^ mice caused a 2-3 fold increase in *Ikbkb* expression at W8 (Fig 2B). Knockdown of *Ikbkb* expression decreased IKKβ protein levels by a comparable 50-60% (Fig 1C) while overexpression of constitutively active *Ikbkb* increased IKKβ protein levels by 2-3 fold (Fig 2D). This deletion efficiency measured at the whole muscle level is comparable to other tamoxifen-inducible models in which satellite and interstitial cells also express the gene of interest. While the deletion efficiency in muscle fibers is likely higher than what is measured at the whole muscle level, it is likely incomplete and thus this model is referred to as knockdown rather than knockout. In male WT mice of the IKKb^MKD^ cohort, *Ikbkb* expression significantly increased by 40-50% in response to tenotomy at both W1 and W8 post-tenotomy (Fig 2A). However, this effect was not seen in the WT mice of the IKKb^MCA^ cohort (Fig 2B), nor was it evident at the protein level (Fig 2C). To ensure that a 50-60% reduction IKKβ protein was sufficient to impact NFκB signaling, we isolated the nuclear fraction from IKKb^MKD^ and WT muscle and quantified NFκB subunits p50 and p65, which reflect the NFκB-proteasomal degradation axis (Kandarian & Jackman, 2006). Nuclear p50 and p65 were reduced by more than 50% in IKKb^MKD^ mice in both sham and tenotomy groups (Fig 2E), confirming inhibition of NFκB nuclear translocation. Thus, these genetic manipulations resulted in a consistent 2-3 fold knockdown or overexpression of IKKβ in both sexes.

**Figure 2.**
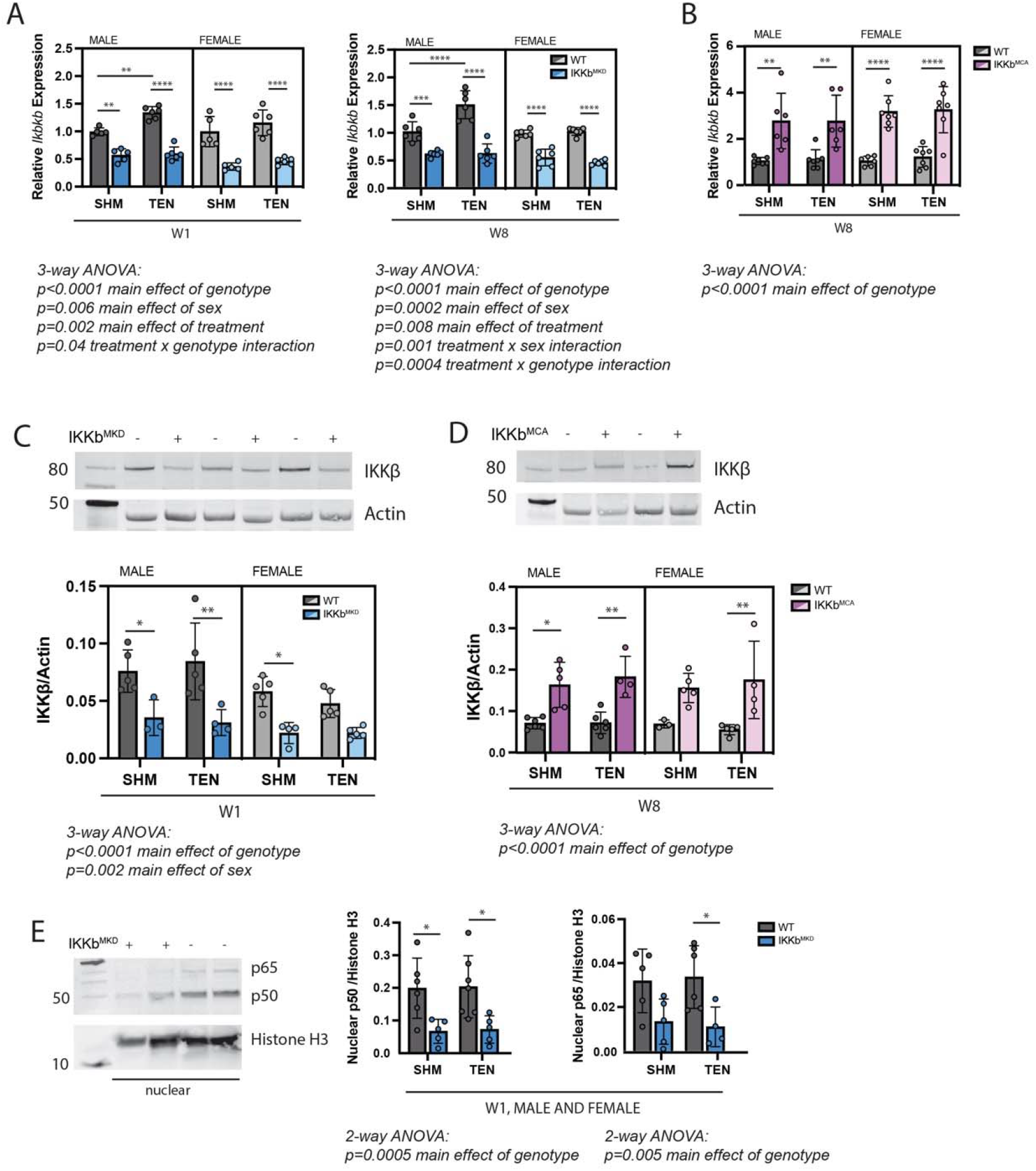
Muscle-specific inducible deletion and overexpression of IKKβ induced a 2-fold decrease and increase in IKKβ expression, respectively. (A) In mice with muscle-specific inducible IKKβ deletion (IKKb^MKD^), expression of the IKKβ gene (*Ikbkb*) was reduced 50-60% in male and female sham (SHM) and tenotomized (TEN) SS at week 1 (W1) and week 8 (W8) compared with wildtype (WT). (B) In mice with muscle-specific inducible overexpression of constitutively-active IKKβ (IKKb^MCA^), *Ikbkb* expression was increased 2-3 fold in SHM and TEN groups at W8. (C) Protein abundance of IKKβ was 50-60% lower in IKKb^MKD^ compared with WT. (D) Protein abundance of IKKβ was 2-3 fold higher in IKKb^MCA^ compared with WT. (E) Protein abundance of NFκB subunits p50 and p65 in the nuclear fraction of SS muscles was reduced 50-60% in IKKb^MKD^ compared with WT. N=3-6 per group; *p<0.05, **p<0.01, ***p<0.005, ****p<0.001.

In both sham and tenotomy groups, IKKβ conditional deletion caused a small increase in mass of the SS (Fig 3A) and IS (Fig 3B) muscles normalized to body mass. 3-way ANOVA applied within each treatment group showed a significant main effect of genotype in SS sham, SS tenotomy, and IS sham comparisons and a trend towards a main effect (p=0.07) in IS tenotomy. However, only in the male SS at W1 post-tenotomy was the individual between-genotype comparison significant. To determine whether IKKβ conditional deletion uniquely affected the atrophic process post-tenotomy or simply increased muscle mass generally, tenotomized SS and IS muscle masses were normalized to the mass of the intact tibialis anterior (TA) of the same mouse. In this normalization scheme, no consistent difference was observed between genotypes in either muscle, either sex, or any timepoint. 3-way ANOVA did not show a significant main effect of genotype in either the SS or IS muscles. Conversely, overexpression of caIKKβ exacerbated tenotomy-induced atrophy to a very minor extent in both SS and IS muscles (Fig 3 D&E), which resulted in a significant treatment x genotype interaction effect by 3-way ANOVA. Normalizing to the mass of the TA eliminated these effects (Fig 3F). Tenotomized muscles in IKKb^MKD^ mice exhibited the same sex-specificity in mass loss (Fig 3A&B) as noted for their WT littermates (Figure 1). Tenotomized muscle mass from the IKKb^MCA^ cohort did not show this effect when normalized to body mass, but there was a significant difference in body mass between sham and tenotomized male mice that may have obscured the atrophic sex-specificity. Both SS and IS masses normalized to TA mass showed significant main effects of sex by 2-way ANOVA (Fig 3F).

**Figure 3.**
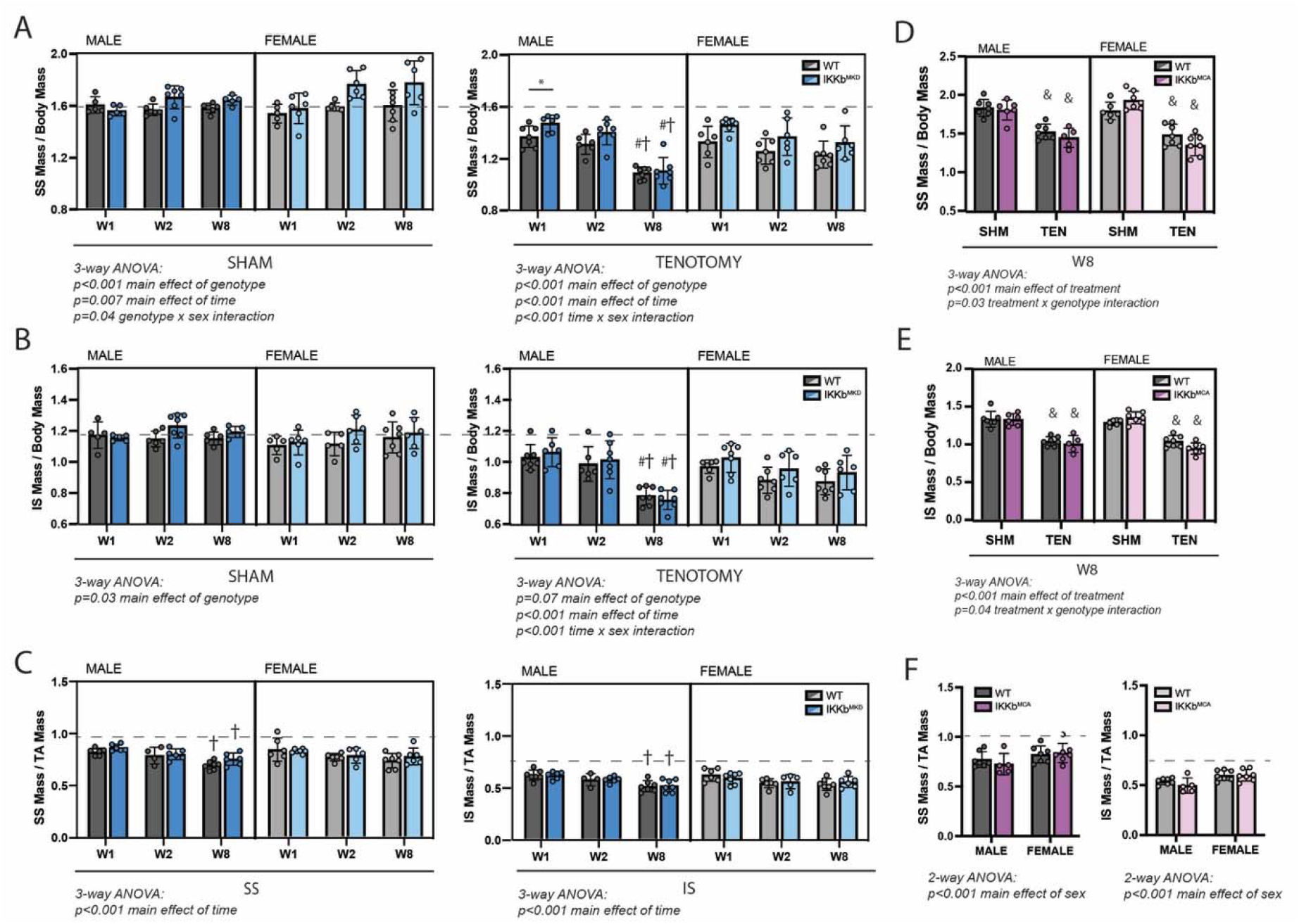
IKKβ deletion and overexpression modestly affected muscle mass in sham and tenotomy groups without uniquely affecting muscle loss post-tenotomy. (A) SS muscles from IKKb^MKD^ mice had a minor increase in mass normalized to body mass in both SHM (left) and TEN (right) groups compared with WT. (B) IS muscles from IKKb^MKD^ mice similarly had a minor increase in normalized mass compared with WT. (C) When SS mass was normalized to tibialis anterior (TA) mass, there was no difference between IKKb^MKD^ and WT groups. (D) IS muscles from IKKb^MCA^ mice had a minor decrease in muscle mass in the TEN group at W8 compared with WT. (E) IS muscles from IKKb^MCA^ mice similarly had a minor decreased in normalized mass compared with WT in the TEN group only. (F) When SS mass was normalized to TA mass, there was no difference between IKKb^MCA^ and WT groups. (A-F) Dotted lines are the average of all sham values included for reference. *p<0.05 between genotypes within the same sex and timepoint. † p<0.05 compared with W1 values within the same sex and genotype. N=5-7 per group; & p<0.05 compared with sham values within the same sex and genotype.

### IKKβ gain- and loss-of-function did not affect tenotomy-induced muscle function loss

Following tenotomy, SS and IS muscles lose ∼20-30% of their specific peak tetanic contractile tension (Fig 4) (Meyer, 2022). This loss was not affected by IKKβ deletion or overexpression of caIKKβ in either muscle or either sex at any timepoint. 3-way ANOVA showed no significant main effects or interaction effects involving genotype. Unlike mass loss, contractile deficits were similar between W2 and W8 in both sexes.

**Figure 4.**
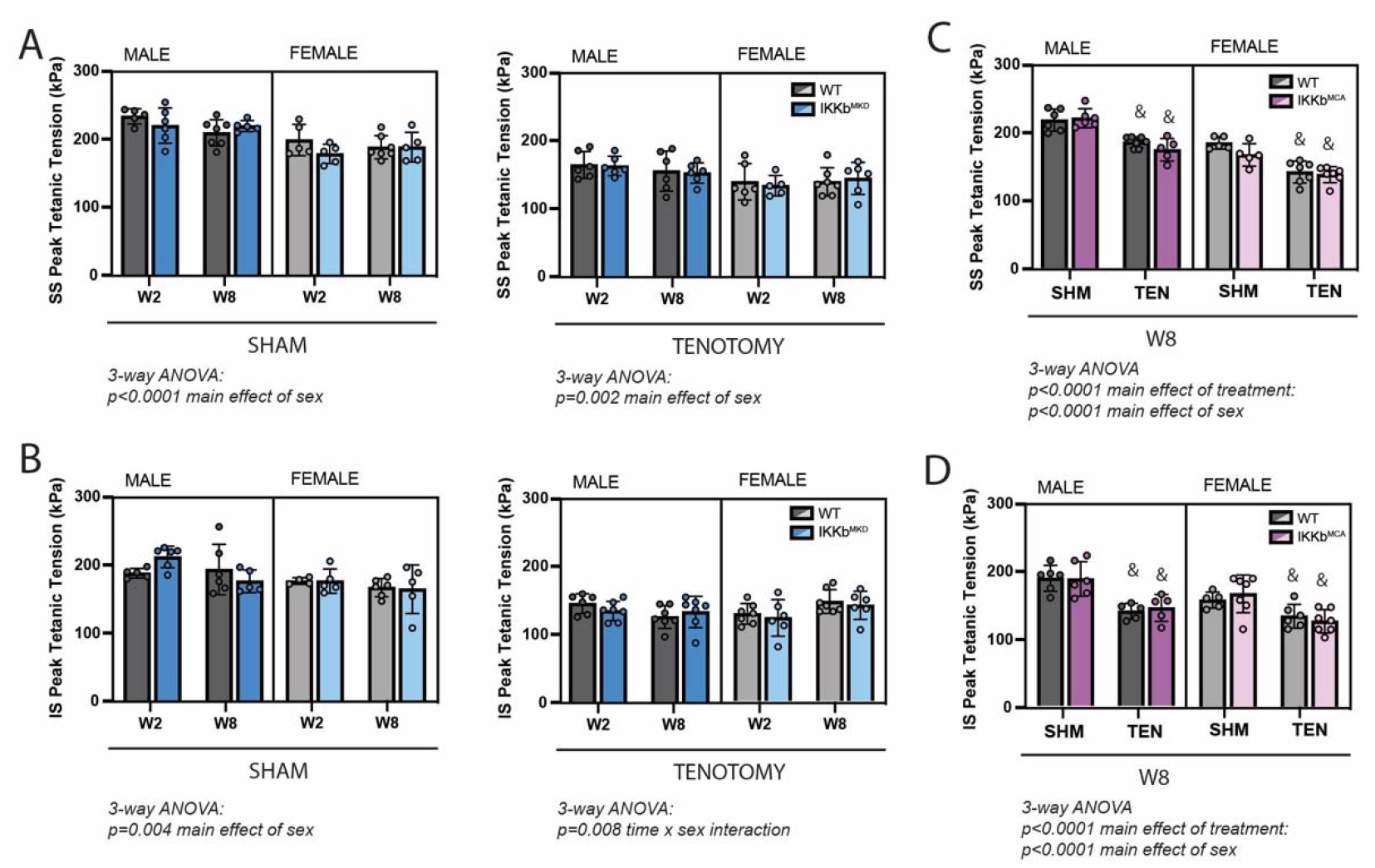
IKKβ deletion and overexpression did not affect the loss of tetanic tension following tenotomy. (A) Peak tetanic tension of SS muscles normalized to physiological cross-sectional area (PCSA) is not different between WT and IKKb^MKD^ genotypes in SHM (left) and TEN (right) groups. (B) Peak tetanic tension of IS muscles is also not different between WT and IKKb^MKD^ genotypes in SHM (left) and TEN (right) groups. (C) Peak tetanic tension of SS muscles is not different between WT and IKKb^MCA^ genotypes in SHM and TEN groups. (D) Peak tetanic tension of IS muscles is not different between WT and IKKb^MCA^ genotypes in SHM and TEN groups. N=5-7 per group; & p<0.05 compared with sham values within the same sex and genotype.

### Architectural adaptations following tenotomy were independent of IKKβ knockdown

We next sought to explore whether the cellular underpinnings of the tenotomy-induced muscle mass loss and contractile dysfunction were modified by IKKβ knockdown. For these assays, we focused solely on the SS muscle, since SS and IS responded similarly to tenotomy with regard to mass and contractile outcomes. Surprisingly, tenotomy caused relatively small changes in fiber CSA compared to other forms of atrophy for a similar mass loss. At W8 (the timepoint of maximal mass deficit), only type 2b fibers (Fig 5A, green) from male muscles exhibited a leftward shift in the histogram of CSA with tenotomy (Fig 5B, green), while type 2a fibers exhibited a rightward shift (Fig 5B, blue). Analysis of mean CSA of type 2a and type 2b fibers over time further illustrated this effect. There was a main effect of time in analysis of type 2a CSA by 3-way ANOVA owing to the continued increase in CSA through W8 (Fig 5C). There was also a main effect of sex, with a significant sex x time interaction effect in type 2b CSA (Fig 5D) owing to the sex-specificity presented in Figure 1. The mass loss unaccounted for by fiber CSA changes was due to decreases in fiber numbers and lengths. ANOVA of fiber numbers (Fig 5E) and fiber lengths (Fig 5F) showed a significant main effect of time owing to their ongoing decreases following tenotomy. Fiber length had an additional main effect of sex (Fig 5F), as the sex-specificity presented in Figure 1 was evident in the IKKb^MKD^ group as well. There were no significant main effects or interactions involving genotype by 3-way ANOVA in any of the comparisons, indicating that IKKβ knockdown did not shift the mechanisms of mass loss. As 3-way ANOVA of type 2a and 2b fiber CSA, fiber number, and predicted fiber length of sham mice showed no significant main effects of genotype or time, average sham values are represented as dotted lines for each sex for comparison (Fig 5 C-F). In contrast to architectural changes, 3-way ANOVA of fiber type distributions showed no significant shifts in any fiber type percentage as a function of tenotomy or IKKβ knockdown (Fig S3).

**Figure 5.**
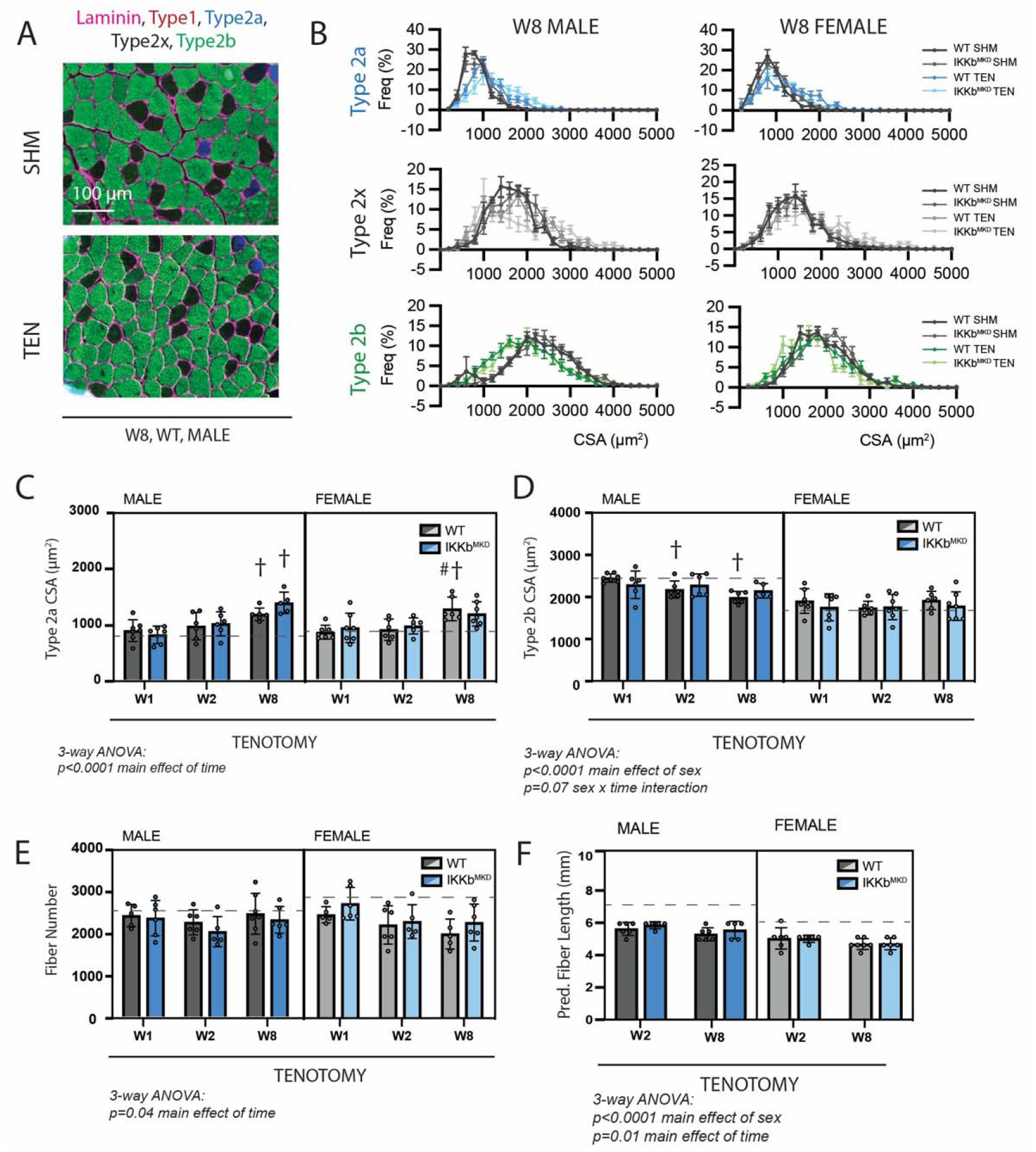
Muscle architectural changes following tenotomy were not affected by IKKβ knockdown and were sex-specific. (A) Representative fiber type staining of histological sections from sham (SHM) and tenotomized (TEN) WT SS muscles at W8. Fiber types were identified by immunostaining against isoforms of myosin heavy chain - type 1 (red), type 2a (blue), type 2x (black), type 2b (green) – and area was quantified by the laminin border (magenta). (B) Distribution of cross-sectional areas (CSA) for type 2a, 2x and 2b fibers for male and female mice at W8. Histograms of type 2b fiber CSA in male mice are shifted to the left following tenotomy. (C) Average CSA of type 2a fibers and (D) average CSA of type 2b fibers across genotypes and timepoints within the tenotomy group. Only type 2b fiber CSA in males decreases following tenotomy. (E) Count of the total number of fibers in the SS cross-section across genotypes and timepoints within the tenotomy group. (F) Prediction of fiber length from measurements of muscle length during physiological testing at W2 and W8 within the tenotomy group. No data exist at week 1 because physiological testing was not performed at that timepoint. (C-F) The dotted lines are the average of all sham values included for reference for each sex. N=5-7 per group; † p<0.05 compared with W1 values within the same sex and genotype.

### Tenotomy-induced structural pathology points to sex-specificity in autophagy

Next, we assessed metrics of structural pathology that are associated with tenotomy and thought to impact muscle contraction – fibrosis, fatty infiltration, and fiber degeneration. Fibrosis and fatty infiltration are progressive in tenotomy (Rubino *et al*., 2007), and were therefore measured at W8 to capture the highest signal. 3-way ANOVA of fibrosis, assessed as the area fraction positive for Sirius Red, showed a significant main effect of treatment, but no main effects of genotype or sex, indicating that tenotomy induced fibrosis equally in WT and IKKβ knockdown males and females (Fig. 6A). 3-way ANOVA of fatty infiltration, assessed as the area fraction positive for Oil Red-O, also showed a significant main effect of treatment, with an additional main effect of sex due to the overall higher levels of intramuscular fat in females (Fig. 6B) (McHale *et al*., 2012). However, there was no main effect of genotype, again indicating that IKKβ knockdown did not affect the progression of fatty infiltration. Fiber degeneration was assessed on H&E-stained sections by a blinded rater counting necrotic fibers (active degeneration) and centrally-nucleated fibers (active regeneration) (Fig. 6C). Across timepoints, there were no obviously necrotic fibers in tenotomized muscle, suggesting that active degeneration was either not occurring or missed at the timepoints chosen for assessment. However, there was a gradual increase in centrally nucleated fibers over time following tenotomy (Fig 6D). 3-way ANOVA of centrally nucleated fibers in tenotomized muscle showed a significant main effect of time owing to this increase and a significant main effect of sex, as this increase was exacerbated in males. The blinded rater noted the appearance of basophilic puncta (hematoxylin positive, DAPI negative structures) in some H&E-stained sections (Fig 6C&E; single arrows). These were generally more prevalent in males than females, and this difference was exacerbated at 8W, where they were increased 20-fold over W1 levels (Fig 6F). 3-way ANOVA of the percentage of fibers with basophilic puncta showed significant main effects of time and sex and a significant time x sex interaction effect. These puncta stained positive for LC3 and p62, suggesting that they were autophagic vesicles.

**Figure 6.**
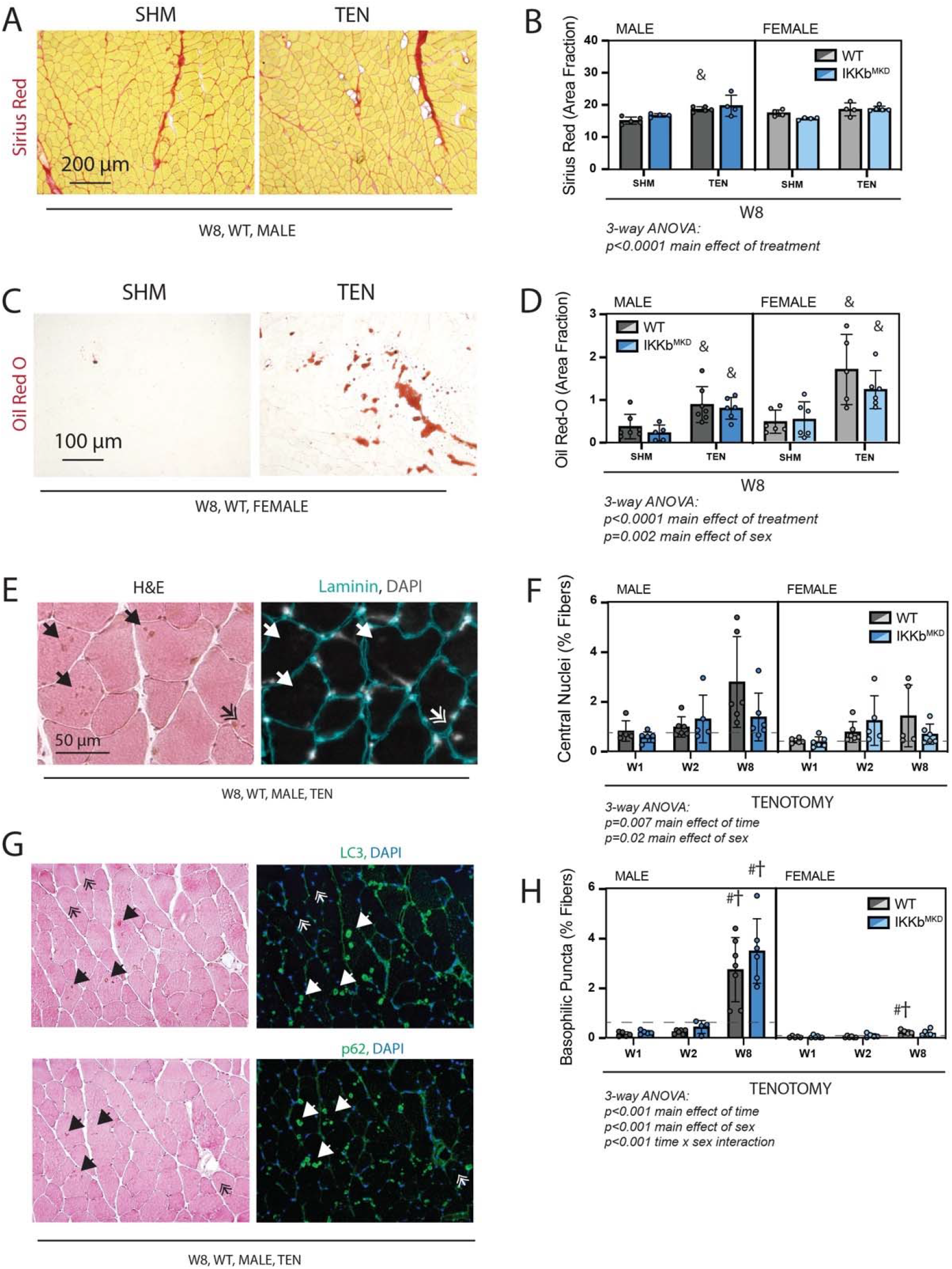
Cellular pathology following tenotomy was not affected by IKKβ knockdown and suggests sex-specificity in autophagic flux. (A) Representative Sirius Red staining of histological sections from sham (SHM) and tenotomized (TEN) WT SS muscles at W8. The red fraction indicates concentrations of collagen. (B) Quantification of the fraction of the section area occupied by Sirius Red as a marker of fibrosis for male and female mice at W8 shows moderate increases with TEN in both genotypes. (C) Representative Oil Red O staining of histological sections from sham (SHM) and tenotomized (TEN) WT SS muscles at W8. The red fraction indicates intramuscular adipocytes. (D) Quantification of the fraction of section area occupied by Oil Red O as a marker of fatty infiltration for male and female mice at W8 shows significant increases with TEN in both genotypes. (E) Representative sections stained with H&E and laminin with DAPI used to identify centralized nuclei (double arrows) as hematoxylin and DAPI positive structures central within the laminin boundary and other basophilic puncta (arrows) as hematoxylin positive central structures negative for DAPI. (F) Quantification of fibers with centralized nuclei as a percentage of all fibers as a marker of regeneration across genotypes and timepoints within the tenotomy group. (G) Representative sections sequentially stained with LC3 or p62 followed by H&E to identify LC3/p62 and hematoxylin positivity in the same section. (H) Quantification of fibers with basophilic puncta as a percentage of all fibers across genotypes and timepoints within the tenotomy group. Basophilic puncta were more abundant in male SS sections at W8. (F,H) Dotted lines are the average of all sham values included for reference for each sex. N=5-7 per group; † p<0.05 compared with W1, # p<0.05 compared with W2, & p<0.05 compared with sham values within the same sex and genotype.

### Knockdown of IKKβ did not induce the typical atrophic signaling pathways after tenotomy

We investigated which atrophic signaling pathways were active during the early and late phases of tenotomy responses. The ubiquitin-proteasome pathway may be activated during the early phase of tenotomy, consistent with a previous report (Valencia *et al*., 2017). Protein ubiquitination, assessed by western blot against ubiquitin, was modestly increased at W1 post-tenotomy, and 3-way ANOVA of ubiquitin abundance showed a main effect of treatment at W1, but not at W8 (Fig 7A). This effect may have been driven by activity of MuRF1, as there was also a main effect of treatment by 3-way ANOVA of MuRF1 gene expression at W1, but not W8 (Fig 7B). At W8, there was a significant main effect of sex that was not present at W1. Interestingly, there were no changes in MAFbx gene expression at W1, but a significant downregulation at W8 (Fig 7C). 3-way ANOVA showed a significant main effect of treatment and sex with significant differences between SHM and TEN groups in male IKKb^MKD^ and female WT and IKKb^MKD^. To examine autophagy, we assessed the protein abundance of LC3. There were no differences in the LC3II/LC3I ratio at W1, but a modest increase at W8 with tenotomy, where 3-way ANOVA showed a significant main effect of treatment (Fig 7D). Overall, neither the ubiquitin-proteolysis nor autophagy-lysosome pathway were substantially affected by tenotomy at W1 or W8. Furthermore, knockdown of IKKβ did not impact these small effects. As the loss of muscle mass could be driven by decreased protein synthesis instead of increased protein degradation, we also assessed activation of the Akt/mTOR pathway, which can mediate increased protein synthesis in response to loading changes. We did not find any changes in phosphorylation of Akt, mTOR, or S6 ribosomal protein with tenotomy or IKKβ knockdown (Fig S2). Finally, there were no differences in myostatin expression, suggesting tenotomy-induced atrophy is also not likely driven by myostatin (Fig S2).

**Figure 7.**
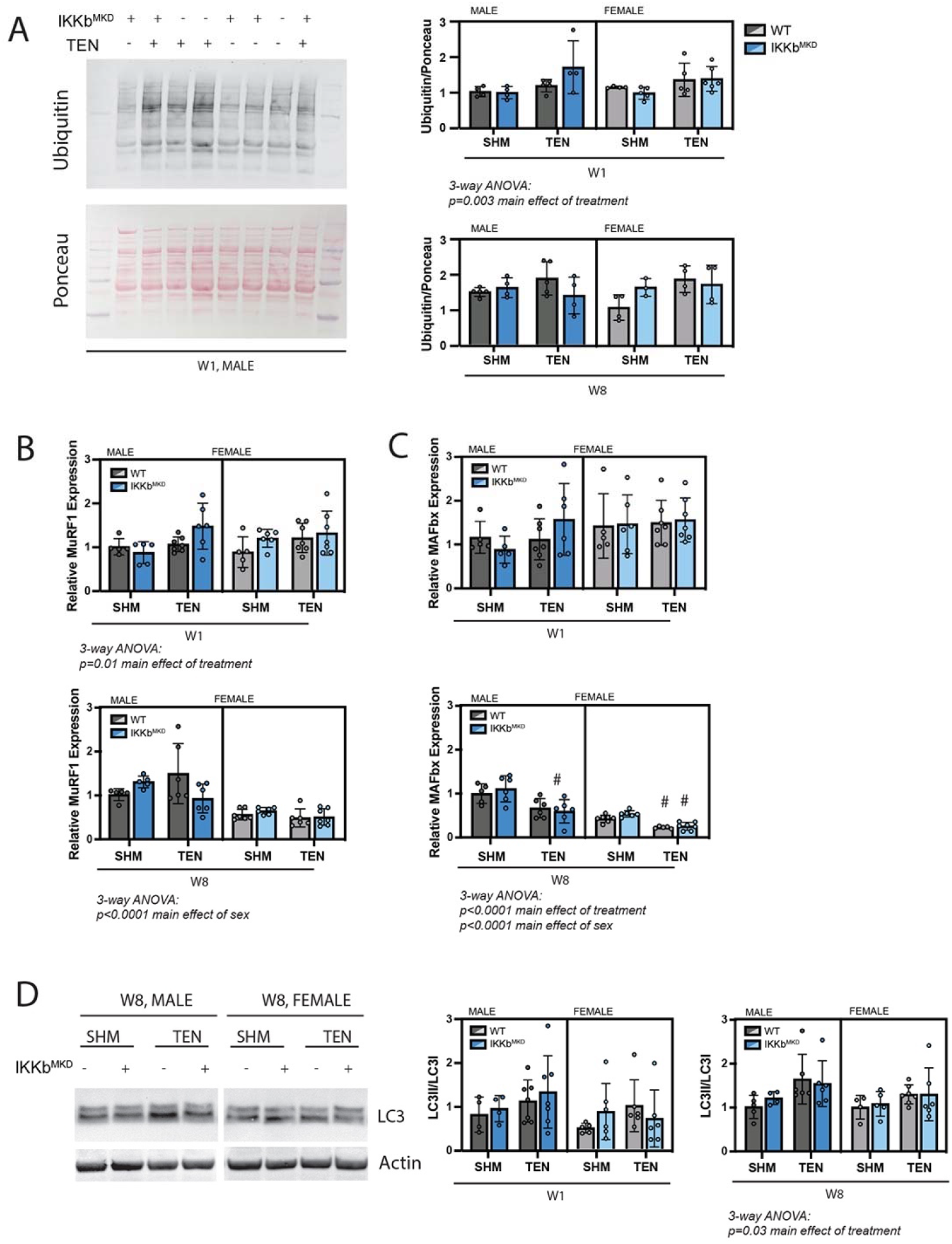
IKKβ knockdown did not impact the major signaling markers of the ubiquitin-proteasome or autophagy-lysosome pathways. (A) Quantification of the abundance of ubiquitin protein normalized to total protein stained with Ponceau at W1 and W8 in male and female mice. (B) Quantification of MuRF1 gene expression at W1 and W8. (C) Quantification of MAFbx gene expression at W1 and W8. (D) Quantification of the ratio of LC3II to LC3I protein abundance at W1 and W8. N=4-6 per group; # p<0.05 compared with W2 values within the same sex and genotype.

## 3 Discussion

This study investigated the effects of NFκB inhibition on muscle atrophy following tenotomy. Using a transgenic muscle-specific and inducible deletion of IKKβ, we were able to block more than 50% of the nuclear translocation of NFκB subunits in muscle fibers at the time of tenotomy of the rotator cuff muscles. While this caused a mild increase in muscle mass in general, it surprisingly had no additional effect on any outcomes following tenotomy, including muscle mass and contractile force, fiber area, length and number and atrophic signaling through the ubiquitin-proteasome and autophagy pathways. Similarly, doubling IKKβ expression using a gain-of-function mouse model had little effect on muscle responses to tenotomy. Together, these results indicate that NFκB does not play a major role in mediating tenotomy-induced atrophy. Interestingly, this study also uncovered sex-specific mechanisms of tenotomy-induced atrophy, most evident at the later timepoints investigated. Male, but not female, mice continued to lose muscle mass between 2W and 8W post-tenotomy. The sex-specific result was likely due to ongoing type 2b fiber atrophy driven by accumulation of autophagic vescicles, which was exacerbated in males at 8W. This surprising finding warrants further investigation in naïve wildtype mice.

The finding that IKKβ knockdown had little effect on tenotomy-induced muscle atrophy aligns with the limited data in tenotomy models, but is contrary to the NFκB muscle atrophy dogma. Transgenic interference with NFκB signaling substantially reduces loss of muscle mass across nearly every model investigated, including unloading, immobilization, denervation, acute inflammatory challenge, nutrient deprivation, and cancer (Cai *et al*., 2004; Mourkioti *et al*., 2006; Judge *et al*., 2007; Van Gammeren *et al*., 2009; Reed *et al*., 2011; Haegens *et al*., 2012; Langen *et al*., 2012; Lee & Goldberg, 2015). These models encompass diverse stimuli for atrophy which are frequently sub-categorized as disuse, aging, or disease (Romanick *et al*., 2013). Tenotomy is typically included in the disuse category along with unloading, immobilization, and denervation. However, tenotomy is fundamentally unique in that it severs the connection of the muscle to the skeleton, eliminating passive tension and immediately changing the length of the muscle as the muscle recoils and comes to rest in a shortened position. Over time, particularly in rodent models, a fibrous “pseudo-tendon” scar develops in the gap between the tendon and bone, partially restoring passive loading to the muscle (Jamali *et al*., 2000). This retraction plus scarring in the shortened position induces the loss of muscle mass in the longitudinal dimension by sarcomere subtraction (Ward *et al*., 2010; Meyer, 2022); a phenomenon which does not occur in hindlimb unloading or denervation models and, to a lesser extent, in immobilization models (Heslinga & Huijing, 1993; Van Dyke *et al*., 2012). Tenotomy and immobilization are also the only disuse models to demonstrate central core lesions of myofibril breakdown, hypothesized to be due to contraction in the shortened position. However, these are also exacerbated in tenotomy compared with immobilization (Baewer *et al*., 2004; Baewer *et al*., 2008). This suggests that tenotomized muscle is engaged in high levels of active remodeling, differentiating it from models reducing excess muscle mass in response to decreased use. This active remodeling could be responsible for the unique degradation signature in tenotomy compared with immobilization or denervation (Bialek *et al*., 2011; Liu *et al*., 2012; Joshi *et al*., 2014), where either the myofibril breakdown or selective serial sarcomere subtraction is engaging autophagic mechanisms. Our data also suggest that the mechanisms of muscle loss may vary temporally following tenotomy. IKKβ knockdown had the greatest effect on tenotomized muscle mass at W1 (Fig 3A), which aligned with main effects of treatment in ubiquitin protein abundance and MuRF1 expression (Fig 7A-B). These transient effects could be due to a differential atrophic responses of the muscle pre- and post-scarring or due to the resolution of inflammatory signals following tenotomy.

Interestingly, while male mice continued to lose muscle mass between W2 and W8 post-tenotomy, the contractile deficit stabilized (Fig 3). Peak tetanic tension is normalized to PCSA, which theoretically accounts for changes in fiber CSA/number and serial sarcomere subtraction, but even after this normalization, a deficit remained. The source of this deficit is unknown. It could be that PCSA is not fully accounting for the impact of architectural changes on contractile force, or the deficit may be attributed to another factor, such as muscle quality. The fact that contractile properties remained stable during ongoing atrophy in male mice suggests that PCSA does sufficiently account for these changes. Additionally, data in the tenotomized rat SS indicate that contractile deficits precede architectural changes and are measurable as soon as one day post-tenotomy, remaining stable for 2 weeks (Valencia *et al*., 2017). This suggests that the contractile deficit originated in the tenotomy injury and never resolved. There is considerable evidence for myofibrillar disruption and fiber damage post tenotomy (reviewed in (Jamali *et al*., 2000)), and it is possible that the tenotomized muscle cannot adequately repair this damage in its unloaded state. If this is the case, it also offers an explanation for why IKKβ knockdown did not affect the contractile deficits.

Expression of a constitutively active form of IKKβ doubled the concentration of IKKβ in mice, yet this increase had negligible effects on tenotomy-induced atrophy. This finding is in contrast to findings from the MIKK mouse and transfection of constitutively active IKKβ, which reportedly induced significant muscle atrophy, even in absence of an unloading stimulus (Cai *et al*., 2004; Van Gammeren *et al*., 2009). However, these studies notable induced >10-fold increases in constitutively active IKKβ, levels which may be supra-physiologic. Our findings suggest that smaller increases in IKKβ may not be sufficient to drive atrophy without an additional stimulus.

Increased protein ubiquitination has been observed early (2-15 days) post-tenotomy (Valencia *et al*., 2017), which could indicate coordinated action of multiple activated pathways, one of which may be regulated by NFκB. Due to this mounting evidence for autophagy due to tenotomy, it was reasonable to hypothesize in the current study that NFκB might also be involved in the atrophic process. Additionally, NFκB could directly affect autophagy as several autophagic regulatory genes (Beclin 1, Atg6, and LC3) are targets of NFκB in other tissues (Copetti *et al*., 2009; Nivon *et al*., 2009). While our data strongly suggest that NFκB does *not* play a major role in tenotomy-induced atrophy, there are several limitations that could have masked some NFκB-driven effects. First, HSA-driven deletion of *Ikbkb* was only 50-60% efficient, leaving 40-50% of the endogenous levels of IKKβ to participate in NFκB signaling. And indeed, we found 40-50% of endogenous WT NFκB subunits p50 and p65 present in the nuclear fraction. By comparison, constitutive deletion of *Ikbkb* under control of muscle creatine kinase (MCK) was 70% efficient (Mourkioti *et al*., 2006), and blocking IKKβ activity using the IKβα super-repressor mouse or transfection of dominant-negative IKKβ completely blocked NFκB nuclear translocation (Cai *et al*., 2004; Van Gammeren *et al*., 2009). The reasons for our limited efficacy are unclear, but may derive from the efficiency of tamoxifen, as ours was the only study to use an inducible transgenic model. However, our lesser impacts on the NFκB pathway are likely more translationally relevant, as NFκB inhibitors have a comparable impact on the NFκB pathway as reported here (Belova *et al*., 2017) and have detrimental off-target effects at high doses (reviewed in (Mourkioti & Rosenthal, 2008)). Thus, while it is possible that residual activation of NFκB signaling is sufficient to drive tenotomy-induced atrophy in our model, our data still suggest that pharmacological inhibition of NFκB signaling is not likely to be therapeutically valuable. Furthermore, our findings in wildtype mice that IKKβ protein levels and nuclear NFκB subunits were unchanged by tenotomy provides additional evidence against NFκB driving tenotomy-induced muscle loss, as both of these are increased in other models of unloading (Hunter *et al*., 2002; Van Gammeren *et al*., 2009). Furthermore, we also found only a very small increase in the major gene target of NFκB-induced atrophy (MuRF1) following tenotomy, consistent with other reports (Bialek *et al*., 2011; Liu *et al*., 2012).

A limitation to this study is that it only examined three timepoints during tenotomy-induced atrophy, the earliest of which was at 1 week post-tenotomy. Inflammation in the rotator cuff is expected to peak early following tenotomy, so it is possible that we missed inflammation-induced NFκB signaling. However, we did not see any durable effects of modifying this signaling, if it exists.
Furthermore, NFκB signaling was active in the tendon at later timepoints in a similar model of tenotomy with repair (Abraham *et al*., 2019). Finally, it is possible that NFκB-driven atrophy was affected by IKKβ knockdown, but compensation by another atrophic pathway obscured the effects on muscle mass etc. Markers of general ubiquitin-proteasomal degradation, autophagy, Akt-mTOR activity and myostatin signaling were not changed by IKKβ knockdown, but this list is not exhaustive.

Few studies have investigated sex-specific effects in muscle atrophy, particularly in the context of tenotomy. Studies in other disuse models indicated that atrophy, and the pathways that modulate it, may vary by sex. Soleus atrophy during hindlimb unloading was exacerbated in female rats compared with male rats (Yoshihara *et al*., 2019; Rosa-Caldwell *et al*., 2021; Yoshihara *et al*., 2022). This was associated with increased FoxO3a activity, protein ubiquitination, and myostatin expression in females compared to males, with no differences in autophagy or Akt phosphorylation (Yoshihara *et al*., 2019). In humans, females experience greater levels of atrophy in response to disuse and aging, while males have greater loss of muscle in response to cancer (reviewed in (Rosa-Caldwell & Greene, 2019)), suggesting that sex specificity extends to clinical populations as well. In the human rotator cuff, symptomatic tear development and progression are more prevalent in males (Yamamoto *et al*., 2017; Song *et al*., 2022). The reasons for this are not clear, but likely include both intrinsic sex dimorphisms and extrinsic environmental and behavioral factors. Here we find that not only do male mice lose more mass after tenotomy, they lose it uniquely through type 2b fiber atrophy, while female mice lose mass uniquely through fiber hypoplasia. It is possible that both sexes experience fiber hypoplasia, but males are more efficiently regenerating the lost fibers; this speculation would explain the sex-specificity in central nuclei, then the load borne by fewer fibers helps maintain fiber CSA. The most dramatic difference we observed between males and females was the increase in basophilic puncta positive for autophagic vesicle markers p62 and LC3 in male muscles at W8 post-tenotomy compared to female muscles. This suggests that the ongoing mass loss concurrent with their appearance may be mediated by autophagy. However, we found only mild differences in LC3 protein content between male and female muscles at this timepoint. Thus, the issue in the male mice may be dysregulated autophagic flux with accumulation of large autophagic vesicles. Proper assessment of autophagic flux cannot be done on existing samples, as it requires treatment of mice with lysomotropic agents prior to muscle harvest. Future studies will examine the interaction between sex, tenotomy, and autophagy in naïve wildtype mice.

In conclusion, we found that a 2-fold gain- or loss-of-function of IKKβ did not impact tenotomy-induced muscle atrophy or contractile dysfunction. This suggests that IKKβ/NFκB inhibitors are unlikely to improve muscle outcomes in human rotator cuff tears. Given promising pre-clinical data suggesting IKKβ inhibitors improve tendon-to-bone healing after rotator cuff tenotomy and repair (Golman *et al*., 2021), and NFκB inhibitors improve muscle mass and contractile function in disease models (Messina *et al*., 2006), it was logical to hypothesize that they would also improve muscle outcomes in human rotator cuff disease. However, this hypothesis was not supported by our data, which suggest that tenotomy, and likely human rotator cuff tears, is a unique model of muscle atrophy driven by factors associated with the loss of muscle passive tension. However, our data also uncovered an intriguing sex-specificity in the atrophic response following tenotomy that warrants further exploration. This aligns with sex-specificity in human rotator cuff tears and could guide more effective and individualized therapies.

## 4 Materials and Methods

### Study approval

All animal work described was performed in accordance with the National Institutes of Health’s Guide for the Use and Care of Laboratory Animals and was approved by the Animal Studies Committee of the Washington University School of Medicine (IACUC 20-0459).

### Experimental design

Experiments were performed on male and female muscle-specific conditional IKKβ knockdown mice (IKKb^MKD^), muscle-specific conditionally active IKKβ mice (IKKb^MCA^) and respective littermate controls at 6-8 months of age. IKKβ-MKD mice were generated by breeding Ikbkbtm2Cgn (originally generated by Manolis Pasparakis) mice with HSA-MCM (Jackson Labs; #025750) mice such that tamoxifen delivery induced deletion of the IKKβ gene in mature muscle fibers. IKKβ-CA mice were generated by breeding Gt(Rosa26)tm4(Ikbkb)Rsky (Jackson Labs; #008242) mice with HSA-MCM mice such that tamoxifen delivery induced expression of a constitutively active form of IKKβ in mature muscle fibers. Littermate mice which did not express Cre-recombinase were used as wildtype (WT) controls. Some data from WT mice have been reported previously (Meyer, 2022). Seven mice per group were randomly assigned in each design - IKKb^MKD^ (3 timepoints, 2 treatments, 2 genotypes, 2 sexes) and IKKb^MCA^ (1 timepoint, 2 treatments, 2 genotypes, 2 sexes) – for a total of 224 mice. Numbers per group for each outcome measure are provided for each figure and deviate from 7 in cases where muscles failed during testing, sample quality was poor or insufficient sample remained for the assay. Each mouse underwent tamoxifen treatment, bilateral tenotomy or sham surgery followed by ex-vivo muscle contractile testing and sacrifice. Muscles were then processed for additional measurements as described below: the SS from the right shoulder and the IS from the left shoulder were used for physiological testing and the SS from the left shoulder and the IS from the right shoulder were prepared for histology, qPCR and western blotting.

### Surgical treatment

Under continuous anesthesia (2% inhaled isoflurane at 2 L/min), both shoulders were prepared for sterile surgery. In a left lateral decubitus position, a ∼3mm incision was made through the skin from the acromial arch toward the mid-belly of the deltoid muscle, exposing the proximal deltoid. An incision was then made through the deltoid to expose the humerus. The humerus was then stabilized with one set of forceps while the acromial space was expanded with another forceps. Mice in the tenotomy group then had the tendons of the supraspinatus (SS) and infraspinatus (IS) muscles transected with microscissors. Mice in the sham group had no tendon transection. All mice then had the deltoid sutured to the trapezius muscle over the acromial arch with 5-0 absorbable Vicryl suture (Ethicon) and the skin incision closed with Vetbond suture glue (3M). The mice were then turned to lie on the right side and the same procedure was repeated for a bilateral treatment. This design was chosen to avoid the effects of altered use that could confound a contralateral control in a unilateral design. Following surgery, mice were provided analgesia and allowed free cage activity until the experimental end point.

### Ex-vivo muscle contractile testing

Mice assigned to the 2 and 8 week post-tenotomy outcome groups were anesthetized again at the experimental end point for ex-vivo contractile testing of the SS and IS muscles as previously described (JOR and JCSM). The SS muscle was excised from the right side with its connection to the scapula and humeral head left intact. All other musculature was dissected free from the scapula, including the IS which was prepared for histology as described below. The SS was then transferred to an ex-vivo physiology rig (Aurora Scientific; 1300A) where it was immersed in Mammalian Ringers solution with the scapula secured to the arm of a dual force/length transducer (Aurora Scientific; 305C-LR) and the humeral head secured to a rigid post. Contraction was elicited by stimulation through parallel plate electrodes flanking the muscle. Muscle length was increased incrementally until the force of twitch contractions began to plateau. Then optimal length was determined by further adjusting length until peak tetanic tension was achieved. Muscle optimal length was then measured with a flexible ruler and the muscle was dissected from bony attachments, blotted, weighed and flash frozen for qPCR. All forces were normalized to physiological cross-sectional area which was calculated from model-predicted fiber length and pennation angle values (Meyer, 2022). A similar procedure was then repeated on the left side for contractile testing of the IS and histological prep of the SS. Following testing, mice were euthanized. Mice assigned to the 1 week post-tenotomy outcome group were euthanized without contractile testing as the connective tissue “pseudo-tendon” was too weak to sustain the connection to the humeral head during testing. Muscles that would have undergone contractile testing were flash frozen in liquid nitrogen and the other muscles were prepared for histology as they were for the other groups.

### Histological measurements

Muscles prepared for histology were affixed to cork at the distal end with tragacanth gum and frozen in liquid nitrogen cooled isopentane and stored at −80ºC until sectioning. They were then transferred to a cryostat (Leica; CM1950) where they were transected at mid-belly with a razor blade. The proximal tip was stored at -80ºC for western blot and qPCR assays. The remainder was then sectioned at the mid-belly face at 10μm for hematoxylin & eosin (H&E), picrosirius red (Sirius Red) and Oil Red-O (ORO) staining and immunostaining for fiber typing by myosin heavy chain isoform (Developmental Studies Hybridoma Bank; BA-F8, SC-71, BF-F3; 1:30), fiber counting by laminin (abcam; 11575; 1:400) outline and autophagic vesicle identification by p62 (Cell Signaling Technology (CST); 5114; 1:100) and LC3A/B (CST; 12741; 1:100).

Sections for ORO staining were mounted on pre-chilled slides and vapor-fixed for 48 hours in formaldehyde at -20 ºC. They were then post-fixed by immersion in 37% formaldehyde for 30 min. For staining, slides were transferred to a working solution of ORO consisting of a 3:2 ratio of ORO stock (5 mg/mL in isopropanol) and diH2O for 10 minutes with agitation. They were then rehydrated and rinsed in diH2O for imaging. Quantification of fatty infiltration was performed on 2x images of the entire cross-section. Sections for Sirius Red staining were fixed in acetone for 1 hour, followed by Bouin’s solution for 5 minutes. Slides were rinsed in running distilled water for 10 minutes and then immersed in a freshly-made Sirius Red solution (1 mg/mL Direct Red 80 dissolved in saturated picric acid at 37ºC, cooled to room temperature) for 2 hours. Slides were then placed in 0.01M hydrochloric acid for 5 min, rinsed in distilled water for 1 minute, dehydrated in ethanol, cleared in xylenes and coverslipped. Quantification of fibrosis was performed on 2 10x images that excluded the tendon and averaged. Images of ORO and Sirius Red were thresholded on the red channel by the Huang algorithm in ImageJ (NIH). Fatty infiltration and fibrosis were then quantified as the percentage of total pixels in the cross-section that met the threshold.

Immunostaining for p62 and LC3A/B was performed on sections fixed in 4% paraformaldehyde for 15 minutes. Following imaging of immunofluorescent staining, sections were stained with H&E allowing positive identification of p62/LC3 and hematoxylin in the same section. All other immunostaining was performed on fresh-frozen sections. With the exception of myosin heavy chain stained sections, all immunostaining was counterstained with Hoechst 33342 (DAPI). Fiber area by fiber type was quantified on 4 20x images from defined regions of the superficial and deep portion of the muscle using a semi-automated ImageJ macro. First, fiber ROIs are identified by thresholding the laminin signal using the Huang algorithm followed by selecting outlined regions by the Analyze Particles algorithm with manual deletion/addition of incorrect/missing regions. Then ROIs are typed by overlaying ROIs on channels stained with each myosin heavy chain type sequentially and scored as positive or negative by average signal. Type assignment is checked and errors corrected manually. Fiber number was quantified on 2x laminin stained images of the entire cross-section using the ROI identification portion of the algorithm. Central nuclei were defined as structures positive for hematoxylin and DAPI not adjacent to a laminin boundary and were counted manually through the entire cross-section. Basophillic puncta were defined as structures positive for hematoxylin but negative for DAPI and were counted manually through the entire cross-section.

### Quantitative real-time PCR

RNA was extracted from frozen muscle using a standard Trizol/chloroform extraction protocol. Briefly, muscle was bead homogenized in Trizol using a TissueLyser II (Qiagen; 85300) and RNA extracted by addition of chloroform with centrifugation. RNA was precipitated in 50% isopropanol, washed in 75% ethanol and dissolved in DNAse/RNAse free water. cDNA was generated using the MultiScribe reverse transcription kit (Applied Biosystems; 4368814) according to the manufacturer’s instructions. Transcript copies were detected using a fast SYBR Green PCR master mix (Applied Biosystems; 4385612) with the following primers: *(Fbxo32: F: AACCGGGAGGCCAGCTAAAGAACA, R: TGGGCCTACAGAACAGACAGTGC), MuRF1 (Trim63: F: GAGAACCTGGAGAAGCAGCT, R: CCGCGGTTGGTCCAGTAG), myostatin (Mstn: F: CAGACCCGTCAAGACTCCTACA, R: CAGTGCCTGGGCTCATGTCAAG) and GAPDH (Gapdh: F: TGTGATGGGTGTGAACCACGAGAA, R: GAGCCCTTCCACAATGCCAAAGTT)*. Reactions were run in duplicate on a QuantStudio3 (Applied Biosystems) real-time PCR system. All expression values were normalized to GAPDH and primers were validated against a positive control (Fig S1).

### Western blotting

Protein was extracted from frozen muscle in RIPA buffer supplemented with Complete protease inhibitor (Roche) with bead homogenization using a TissueLyser II (Qiagen; 85300). Homogenized tissue was then solubilized for 1 hour at 4 ºC with agitation, centrifuged and the protein concentration of the supernatant determined by a Pierce BCA assay (Thermo Fisher Scientific; 23225) according to the manufacturer’s instructions. Then, equivalent amounts of protein (40 μg) diluted in diH2O with laemmli buffer were denatured and separated on 4-12% Bis-Tris gels (Invitrogen; NW04120). Protein was then transferred to polyvinylidene difluoride (PVDF) membrane, reversibly stained with ponceau S and blocked in TBST+ (1x Tris-buffered saline with 2.5% fish gelatin, 0.1% sodium azide and 0.5% tween). The following primary antibodies were applied overnight at 4 ºC at 1:1,000 unless otherwise noted: IKKβ (CST; 8943), Ubiquitin (Dako; Z0458), p62 (CST; 5114), LC3A/B (CST; 12741), mTOR (CST; 2983), phospho-mTOR-ser2448 (CST; 5536), Akt (CST; 9272), phospho-Akt-ser473 (CST; 4060) and Actin (Sigma; A2066; 1:10,000). To assay NFκB subunit translocation, protein from a subset of muscles was fractionated into nuclear and cytoplasmic components using a Nuclear Extract Kit (Active Motif; 40010) according to manufacturer’s instructions. The nuclear fraction was validated against an established protocol (22994964) and separated, transferred and blocked as described above. The following primary antibodies were applied overnight at 4 ºC at 1:1,000 unless otherwise noted: p65 (CST; 8242), p50/105 (CST; 13586), Histone H3 (CST; 4499), and GAPDH (abcam; 9485; 1:10,000). Following incubation, membranes were washed and incubated for 1 hour with the relevant secondary antibodies and imaged with a LI-COR Odyssey. Blot analysis was performed using Image Studio (LI-COR). Band intensities were normalized to an actin loading control with the exception of p50 and p65 which were normalized to Histone H3 and Ubiquitin which was normalized to Ponceau. When possible, antibodies were validated with a positive loading control (Fig S1).

### Prediction of mass deficit from architectural measurements

The relative contribution of fiber CSA, length and number changes to the mass deficit was calculated by modeling fibers as cylinders. Under this assumption, the mass loss in a fiber (Δ*m*) resulting from a decrease in fiber length (Δ*FL*) and decrease in fiber CSA (Δ*F*_*CSA*_) was calculated by multiplying the two and the density of muscle, ρ (Ward & Lieber, 2005). Then, the mass loss in the muscle was calculated by multiplying the mass loss in individual fiber types by the relative fraction of each fiber type (Rui *et al*., 2016) and the decrease in fiber number (ΔFN), according to Equation 1. This equation assumes that the density of muscle and the relative fraction of each fiber type is unchanged following tenotomy.

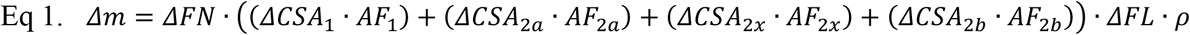

### Statistical analyses and reproducibility

Sample size was selected a priori using a power calculation (G*Power) with variance estimated from measures of SS mass and peak tetanic tension from a previous study. With n=7, this study was predicted to be able to detect a 15% difference in sample means of both variables with α=0.05 and (1-β)=0.8. Following genotyping and non-randomized group assignment, mice were given a numeric identifier and all experimenters were blinded to genotype. Cage-mates were assigned to a single treatment to enable true littermate controls. Between group comparisons were made by 2- or 3-way ANOVA as indicated, with Sidak’s or FDR multiple testing correction applied, respectively. Normality was tested using the Shapiro-Wilk test. Actual numbers per group are indicated in each figure for each analysis. All results are presented as mean +/-standard deviation. All statistical analyses were performed with GraphPad Prism.

## 5 Data Availability

All data generated or analyzed during this study are included. Data files have been provided for all Figures.

## 7 Author Contributions

GAM and ST were responsible for the conception and design of the study. GAM and KCS contributed to acquisition, analysis and interpretation of the data. GAM was responsible for the drafting of the manuscript and GAM, ST and KCS all contributed to critical revision and gave final approval.

## 9 Acknowledgments

Special thanks to Kathryn Bohnert and Heangun Yoon for technical assistance and Dr. Drew Findlay for insightful advice and discussion.

---

**Figure S1.**
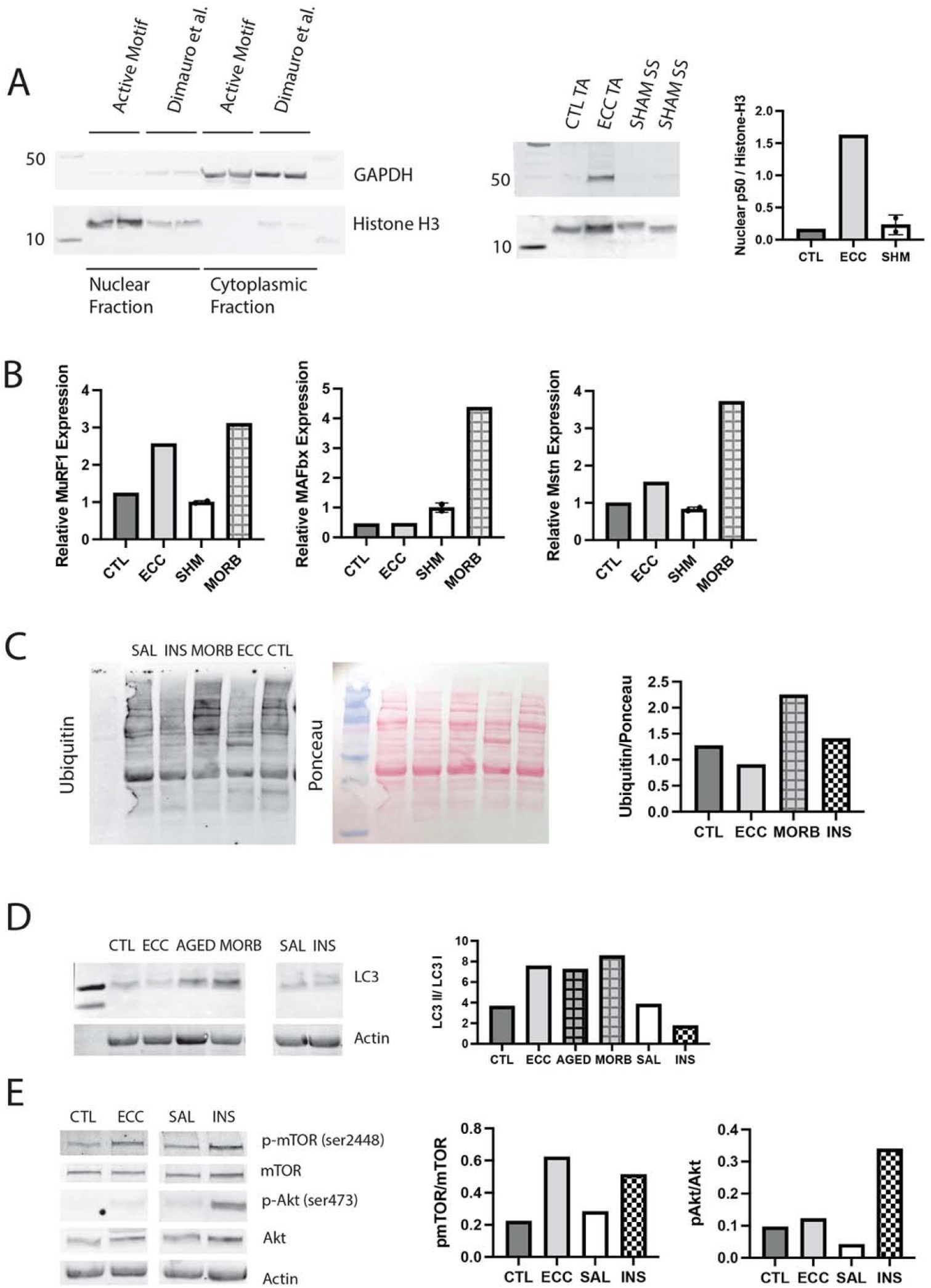
Validation of assays with positive controls. (A) Validation of the Active Motif nuclear extract kit against a literature validated protocol designed for skeletal muscle (left). Western blot of nuclear p50 in protein extracts from a tibialis anterior (TA) muscle subjected to 50 eccentric contractions (ECC) one hour prior to harvest, the contralateral control TA (CTL) and two samples from the W1 WT male sham (SHM) group (right). Nuclear p50 is expected to be elevated following eccentric exercise (Jiménez-Jiménez *et al*., 2008) (B) Expression of MuRF1, MAFbx and myostatin (Mstn) genes in CTL, ECC, SHM and the infraspinatus from a 24 month old moribund mouse (MORB). MuRF1 expression is expected to be elevated following eccentric exercise, while MAFbx remains unchanged and myostatin is decreased (Louis *et al*., 2007). Both MuRF1 and MAFbx are expected to be elevated in critical illness (Wollersheim *et al*., 2014). (C) Western blot of ubiquitin in muscle extracts from CTL, ECC, SHM, MORB and a 24 month age matched healthy IS (AGED). Ubiquitin is expected to be elevated with eccentric exercise and critical illness (Murton *et al*., 2008). (D) Western blot of LC3A/B in muscle extracts from CTL, ECC, AGED, MORB and a mouse treated with insulin (INS) one hour prior to harvest and a saline (SAL) treated control. The ratio of LC3II / LC3I is elevated under increased autophagic flux which is expected with eccentric exercise and critical illness, but not with insulin treatment (Banduseela *et al*., 2013; Langer *et al*., 2021). (E) Western blot for signaling proteins p-mTOR (Ser2448), mTOR, p-Akt (Ser473) and Akt in CTL, ECC, SAL and INS muscle extracts. p-mTOR and p-Akt are expected to be elevated relative to total mTOR and Akt with insulin treatment, but only p-mTOR is expected to elevated following eccentric contractions (Parkington *et al*., 2003).

**Figure S2.**
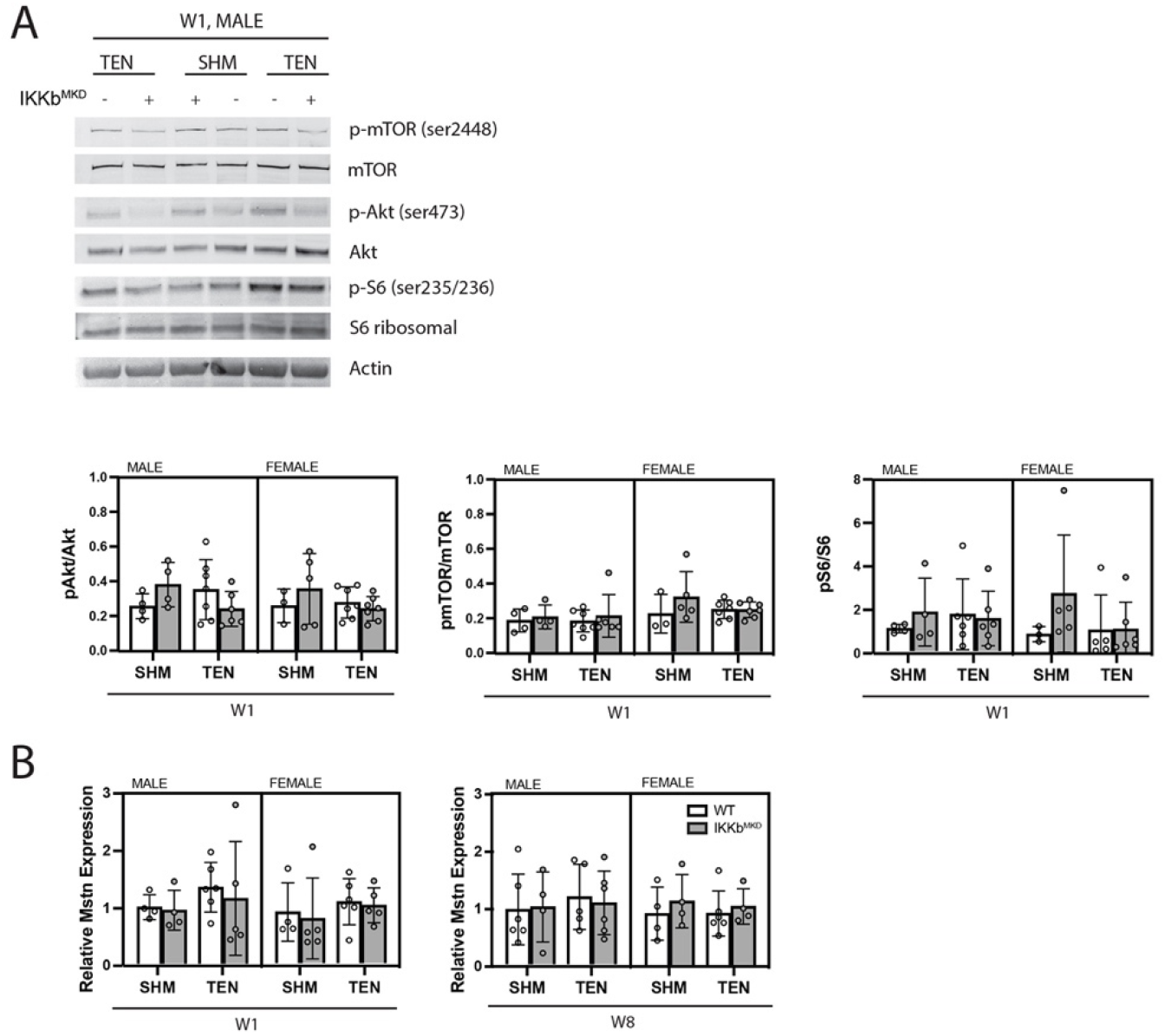
Akt/mTOR/S6 Ribosomal phosphorylation is unchanged in response to tenotomy or IKKβ knockdown. (A) Representative western blot for signaling proteins p-mTOR (Ser2448), mTOR, p-Akt (Ser473), Akt, pS6 ribosomal (ser235/236) and S6 ribosomal for W1 male samples. Quantification of phosphorylated protein abundance relative to total respective protein abundance across treatment, genotype and sex. (B) Gene expression of myostatin (Mstn) at W1 and W8.

**Figure S3.**
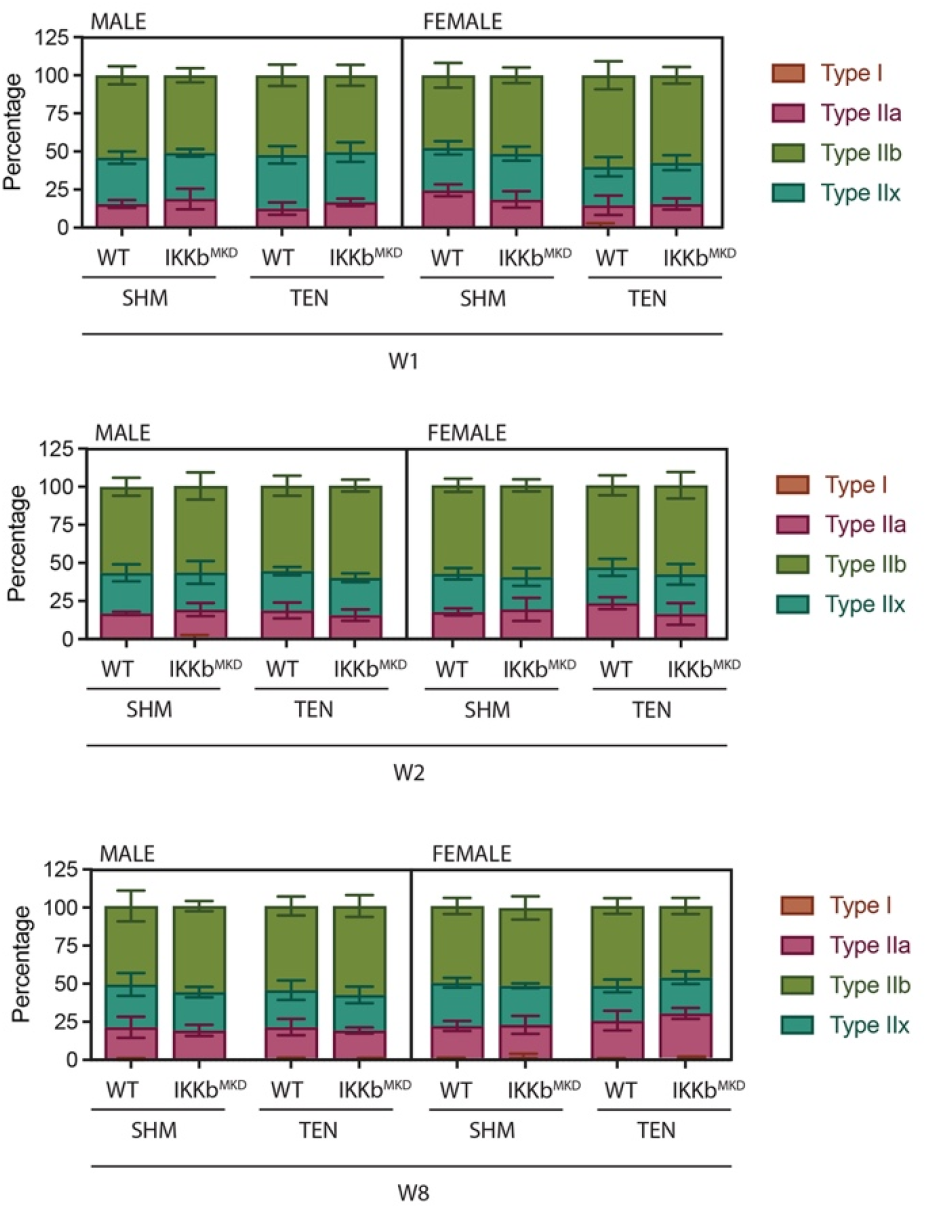
Fiber type distribution is not altered by tenotomy or IKKβ knockdown. Distribution of fiber types as quantified by myosin heavy chain isoform immunostaining on histological sections.

